# Single-Cell Microwave Cytometry for Drug Resistance Detection in Cancer

**DOI:** 10.1101/2025.06.24.661288

**Authors:** Uzay Tefek, Hashim Z. Alhmoud, Tieu Lan Chau, Sayedus Salehin, Gizem Damla Yalcin, Ozge Saatci, Burge Ulukan, Scott Auerbach, Beana Puka, Ahmet Acar, Serkan Ismail Göktuna, Ilyas Chachoua, Ozgur Sahin, M. Selim Hanay

## Abstract

Monitoring biophysical changes in single cells induced by drugs is crucial for advancing cancer therapies, especially for highly heterogeneous tumours. Emerging methods for single-cell drug sensitivity testing have largely relied on phenomenological analyses. Here, we present a novel electronic cytometry platform integrating microwave resonators with impedance cytometry, enabling simultaneous, label-free measurements of cell volume and dielectric permittivity at microwave frequencies. This approach uniquely captures intracellular differential responses indicative of drug-induced biophysical states, thus providing deeper insights beyond traditional phenomenological analyses. We first validated the platform using varying salt concentrations. We then demonstrated that the sensor platform could differentiate between drug resistant versus sensitive phenotypes in multiple isogenic cancer cell lines treated with cytostatic, cytotoxic and mixed-effect drugs. Notably, we showed that the technique performs successfully in patient-derived tumor organoids, a model system highlighting its immediate clinical relevance. Our findings underscore the potential of electronic measurements at the single cell level to provide informative and actionable biophysical signals relating to cellular drug response. The implementation of this platform could significantly advance cancer treatment by identifying the resistant cell populations in heterogenous tumours and optimizing selection of drugs for each patient, paving the way towards precision medicine.

## Introduction

Chemotherapy has been the mainstay therapy for several different cancers with proven success; however, its effectiveness is often limited by the tumour heterogeneity and the development of drug resistance. In contrast, targeted therapies directed towards the drivers of carcinogenesis are more effective and have fewer side effects, as they are designed to interfere directly with the specific molecular mechanisms responsible for tumour development and progression while keeping normal cells unaffected.^1^ Consequently, patients often experience better outcomes, less toxicity, and an improved quality of life. Unfortunately, patients eligible for targeted therapies represent only a small fraction of the affected population.^2^ Moreover, even among personalized therapies that reach the clinical setting,^3^ only a small subset of patients respond effectively,^4,5^ while many of those who respond to the treatment are likely to develop drug resistance after initial remission.^2^

To address these challenges, precision oncology uses a multitude of tools to predict a patient’s response to therapy, including DNA and RNA sequencing, protein expression profiling, and multiomics-based biomarker detection. However, there are certain challenges involved in these approaches. For example, while the identification of single genes has yielded promising clinical results, a more in-depth view of clinical cases often reveals complex networks of interacting gene expressions that ultimately affect cancer development and metastasis. On the other hand, research into high-throughput multi-omics has yielded enormous volumes of data which are not easy to translate into disease phenotypes clinically.^1,6^ Therefore, there remains an unmet need to create rapid and effective strategies to identify the emergence of drug resistance early on in tumours having high heterogeneity.

An emerging new approach is functional precision medicine where the effectiveness of specific drugs are directly tested on living cells isolated from patient biopsies and their efficacy immediately evaluated using *in vitro* standard viability assays.^2,7^ However, functional precision medicine encountered limitations in wide-scale use due to large sample requirements, long turn-around times, genetic drift in isolated samples, and the loss of cell viability *ex vivo*.^8^ The challenges mentioned above can be overcome by the advent of rapid microfluidic-based single-cell biosensors.

In the field of single-cell sensors, monitoring changes in cellular biomass induced by drug exposure emerged as a robust and non-invasive assay for functional precision medicine in recent years.^9,10^ Cellular mass measurements probe cellular composition which is affected by the action of drugs. Cellular mass is measured most sensitively by suspended microchannel resonators (SMRs), where a hollow micro-cantilever with an integrated microfluidic channel carries the media containing single cells.^11–13^ This assembly is oscillated in vacuum to obtain the buoyant mass of each cell passing through the hollow micro-cantilever. In certain cases, SMR technology can detect cell pathology by means of measuring changes in cellular mass even more rapidly than through conventional assays.^14^ This observation, in combination with high-throughput microfluidic-based measurements and low sample volumes allows the development of novel testing modalities with faster turnaround times for informed drug selection.

However, focusing on cellular changes solely in one dimension such as cellular mass — which significantly varies through the cell cycle —coupled with the delicate device and apparatus requirements to accomplish such measurements has so far limited the reach and scope of this approach. Recent advances have highlighted the potential of multi-parametric biophysical measurements for cell density, including high-throughput profiling using fluorescence exclusion microscopy coupled with SMRs.^15^ Here, we take a different approach to probe cellular composition electronically and along multiple dimensions by using a simple microfluidics system which can be fabricated in standard cleanroom facilities. With this system, it is possible to simultaneously obtain cellular volume, electric size (an analogue of dry mass in the electrical domain), and dielectric permittivity. These parameters can then predict the outcome of an anti-cancer drug treatment.

Recently, we have reported the development of an electronic sensor which integrates a low-frequency Coulter counter, and a high frequency microwave resonator to probe single microparticles such as microplastics and single cells.^16^ These sensors simultaneously measure the cellular volume and electrical size. Electrical size measurements operate at microwave frequencies, which bypass ionic contributions and instead detect the dielectric contrast between cellular biomaterial and water — making electrical size an electronic analogue of a cell’s dry mass. By combining the cellular volume and electrical size, it is possible to obtain the microwave permittivity of cells in a high-throughput manner. In this sense, the measurements in the microwave domain yield a metric for the amount of biomaterials and water inside cells, thus providing a unique fingerprint for the material composition of a cell. When the frequency of measurement lies within the dipolar polarization regime (1-10 GHz), microwave measurements yield the electrical analogue of cellular mass and mass density, which can be utilized to monitor the biophysical state of a cell when exposed to drugs (Figure 1).

**Figure 1.**
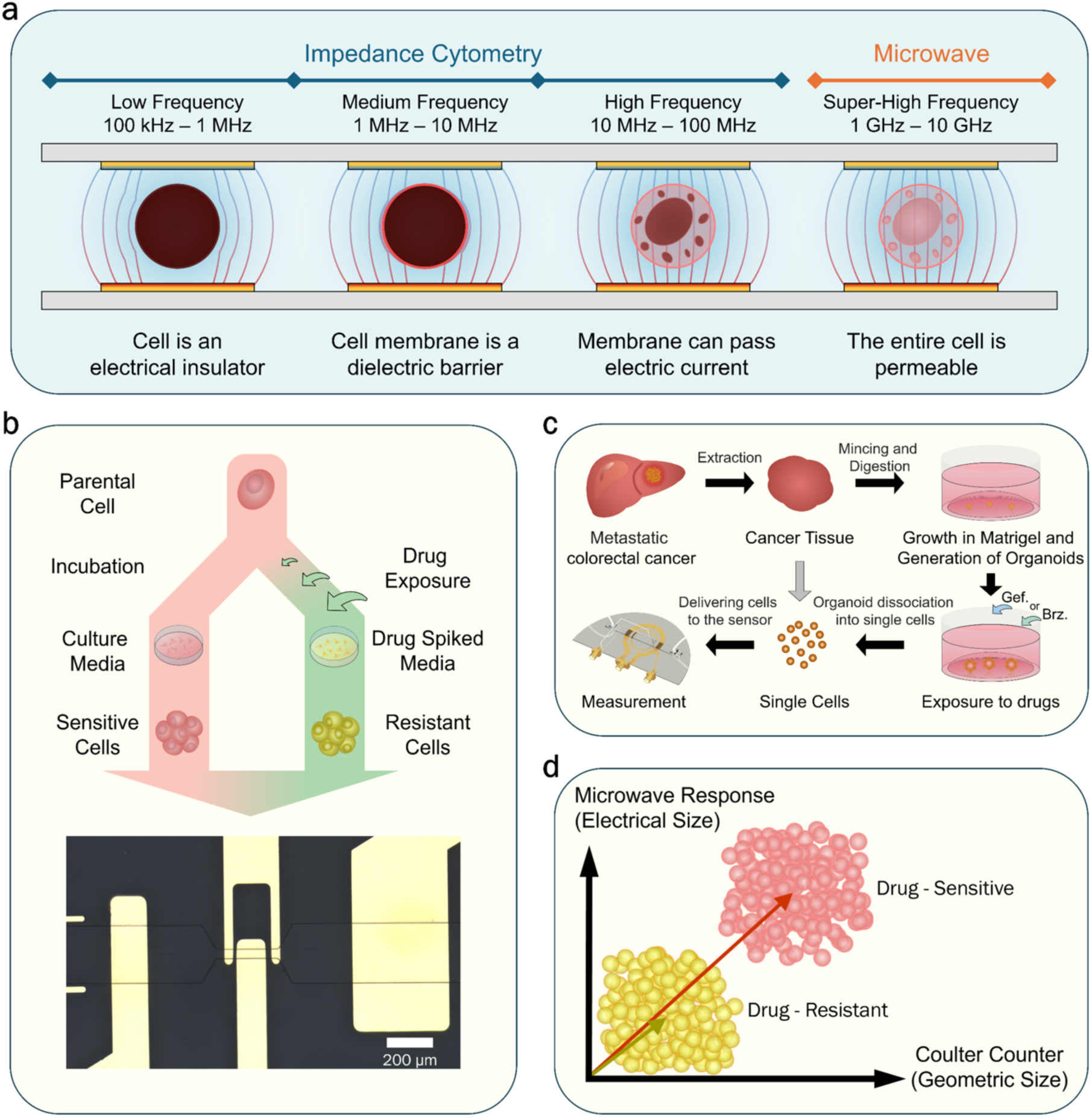
Schematic representation of the sensor operating principles. a) Shows how the increase in the frequency of the alternating electric field ends up penetrating into the cell interior as it no longer suffers from charge-shielding effects of the ions, since ion movement cannot keep up with the speed of the alternating electric field at GHz frequencies. b) Shows the workflow followed to generate and test resistant cancer cell line from their parental counterparts, c) Shows the workflow followed to generate patient derived organoids and measure their susceptibility to Gefitinib (Gef) and Bortezomib (Brz) drugs (black arrows). Grey arrow indicates an alternative, and faster path for testing single cells. d) Indicates how parental cells were differentiated from their resistant variants by looking at the change in the measured permittivity when exposed to the drug.

Indeed, the changes in the biomaterial-to-water ratio inside a cell are better reflected in the microwave permittivity of the cell, rather than the mass density. This is because the difference in electrical polarizability is very large between water (𝜖_𝑟_ ≈ 70) and hydrocarbon chains inside organic groups (𝜖_𝑟_ ≈ 2 − 3); on the other hand, there is not much difference between these two types of components in terms of mass density (1.0 g/cm^3^ vs 1.4 g/cm^3^). Thus, the internal permittivity of a cell is inherently a more sensitive probe into the composition of a cell. Owing to the simplicity of the sensor fabrication, the absence of any high frequency mechanical oscillations and suspended structures with narrow channels, and independence from optical or biochemical labelling, the sensor avoids the pitfalls of state-of-the-art technologies in terms of complexity, high-cost, and the need for expert operators, while maintaining all the benefits of high-throughput microfluidic-based sensors, offering flexibility, sensitivity, and speed. Moreover, the developed platform requires only small sample sizes on the order of a few thousand cells per measurement set.

Here, we demonstrated the capabilities of this sensor by first investigating changes in volume and electrical size of single cells by doing a proof-of-principle experiment where we used hypo- and hypertonic solutions to induce water uptake in human embryonic kidney (HEK293 FT) cells. By combining these measurements, we tracked the changes in cell microwave permittivity (ε) as a function of solution osmolarity at the single-cell level. For drug testing, we began with a human colorectal carcinoma model (DLD-1) and monitored the effects induced by gefitinib (EGFR inhibitor) on both parental and drug-resistant variants using measurements of geometric and electrical size, permittivity, and —for independent verification— Trypan Blue viability assay. We then exposed the same cell populations to a different drug (cisplatin) to verify whether the observed resistance was drug-specific and whether single-cell microwave cytometry can resolve the drug-specific effects. We extended these measurements for two more cell lines (in each case with parental and resistant variants) against cisplatin and tamoxifen. These three drugs represent different modes of action of cancer therapeutics — relatively cystostatic (gefitinib), cytotoxic (cisplatin) and mixed-effect i.e., both cytostatic and cytotoxic (tamoxifen) — allowing us to evaluate sensor’s ability to distinguish diverse biophysical signatures associated with distinct modes of action. Indeed, in all cases, the electronic measurements recapitulated the biological observations about the cellular viability, drug susceptibility, and development of drug resistance accurately. Finally, we tested the sensor with Patient-Derived Organoid (PDO) models established from metastatic colorectal carcinoma from a cancer patient against two drugs with known differential efficacy —bortezomib (effective) and gefitinib (ineffective). Overall, the results demonstrate that cellular drug response can be assessed accurately by this integrated electronic approach through single-cell measurements and at high throughputs, underscoring its potential as a functional precision medicine tool for guiding therapy selection.

## Results

### Probing the effects of hypo- and hypertonicity on cell volume and permittivity

Measuring the microwave permittivity (ε) of the cell interior provides an effective assessment of the ratio between biomaterial and water content of the cell. Biomaterial content encompasses proteins, nucleic acids, carbohydrates, and lipids. The relative permittivity of pure water is approximately 70 at microwave frequencies, whereas proteins exhibit relative permittivity values ranging from 2.5 to 3.5 (bulk values^17^), resulting in a substantial and measurable contrast between these materials. Additionally, the biomaterial-to-water ratio dictates the reaction kinetics inside the cell, by determining the reaction rates and equilibrium concentrations in biochemical processes.^18^ The biomaterial-to-water ratio exhibits itself in cell mass density which is a tightly regulated property of mammalian cells,^18–20^ as variations in enzyme concentration shift the biochemical equilibrium and can affect proper biochemical functions, structural integrity, and protein folding.^21,22^ Notably, only a small number of processes can affect cell mass density, including cell division, differentiation, apoptosis, senescence, and changes in extracellular osmolarity.^18^ On the other hand, protein synthesis and protein degradation appear to have minimal impact on cell density.^23^

Under hypotonic stress, cells uptake water to equalize osmotic pressure across the lipid membrane. This uptake naturally entails a change in the intracellular density. At osmolarities as low as ∼140 mOsm/L (*i.e.*, approximately half the physiological osmolarity), cells generally maintain their nominal volume^24^, and only begin to uptake water at smaller osmolarities. To assess whether these changes in density can be measured by our sensor, we subjected HEK293 FT cells to various concentrations of PBS solution at varying concentrations (between 10% and 150%) and measured the cell volume and electrical size, at the microwave band, approximately one minute after resuspension. Fig. 2a illustrates the changes in the geometric and electrical volume of HEK293 FT cells at varying concentrations (10%, 20%, 50%, 100%, and 150%; 28.8, 57.5,143.8, 287.5, and 431.2 mOsm/L, respectively). Microwave permittivity (ε) was derived by normalizing the electrical diameter to the geometric diameter. Fig. 2b illustrates the changes in permittivity (left-axis) and geometric diameter (right-axis) by Earth Mover’s Distance (EMD) which is a measure of dissimilarity between two statistical distributions.^25^ An interesting effect was observed in geometric volume for diluted concentrations: moving from an isotonic condition to 50% dilution, the change in the geometric diameter was quite small (<10% increase) matching what has been shown in the literature.^24^ However the geometric diameter changed significantly for the 20% and 10% dilutions (+25% and +38% increase respectively).

**Figure 2:**
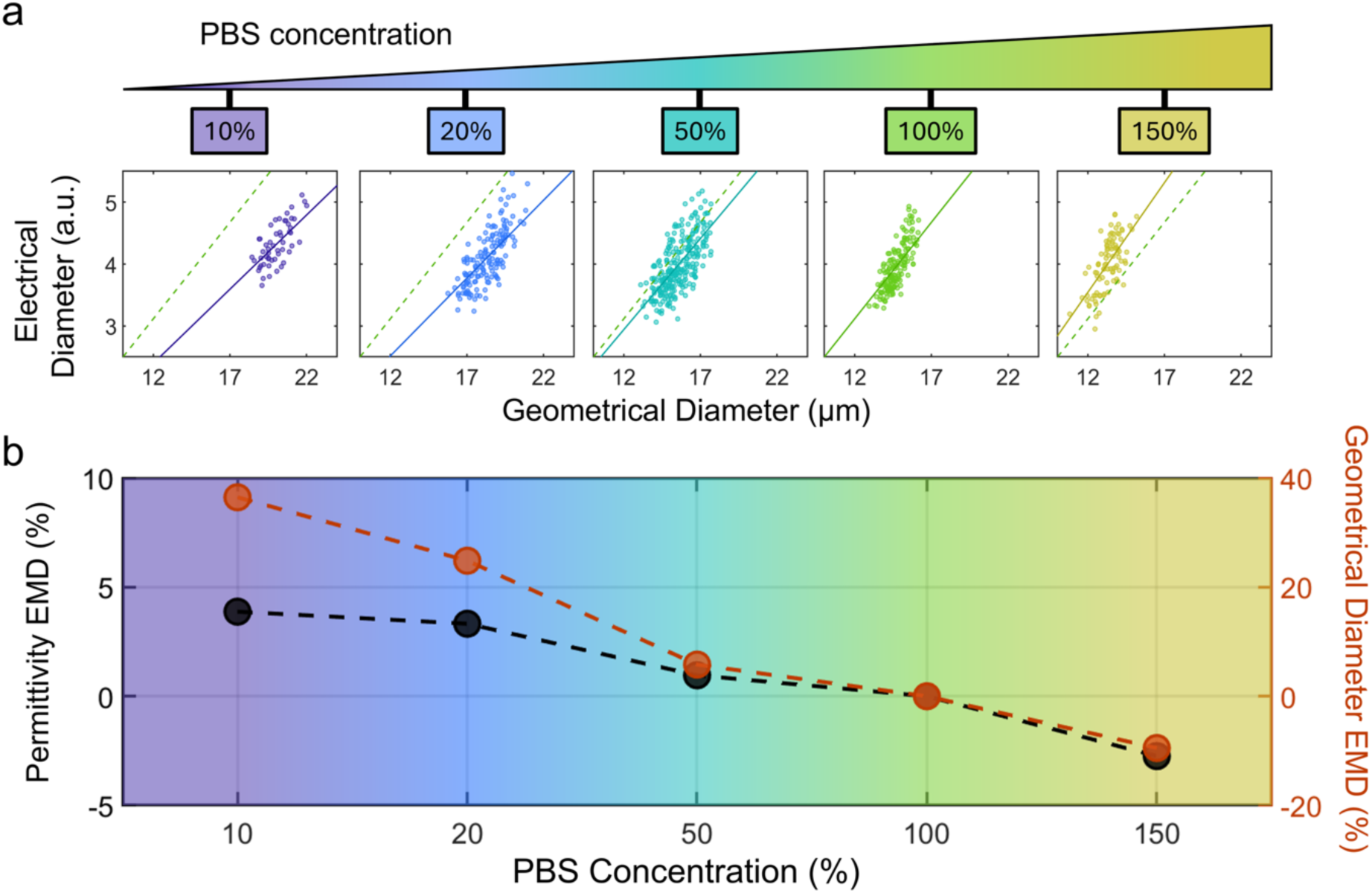
Effect of salt concentration on the cell. a) Shows electrical vs. geometrical diameter change as a function of PBS concentration, where 100% refers to undiluted PBS. b) The change in permittivity and geometric diameter using Earth Mover’s Distance (EMD) expressed as percentage change, as a function of the varying concentration of PBS.

Concurrently, we also observed a change in the electrical permittivity (ε) matching the expansion in cell volume. The change was minimal at 50% dilution (∼1% increase) resulting from the small volumetric increase and uptake of water. This change rose to 3% and 4% for the 20% and 10% dilutions respectively. Normally under hypotonic conditions, cells facilitate the efflux of electrolytes (Na^+^, K^+^, and Cl^-^) to equilibrate the cytosolic osmolarity with the external environment. However, at high (GHz) frequencies, ions in solution do not significantly contribute to polarization (and thus ε) extracted by our sensor, because the ionic current is significantly dampened at GHz frequencies.^26^ Therefore, we hypothesize that the change in ε is due to compositional changes relating to the ratio of biomaterial in the cell, relative to water, starting to decrease with water uptake, as opposed to the ionic concentration decreasing in the cytosol. Previous studies have shown that cells indeed expel organic osmolytes in addition to electrolytes in the early stages of exposure to a hypotonic environment in order to equilibrate osmotic pressure before resorting to volumetric changes.^24,27^ In contrast, at more extreme dilutions, the cells will finally resort to water uptake in order to equalize pressure as seen at 20% and 10% PBS concentrations, but the trend of increasing ε continued as the cell mass density decreased in parallel. Furthermore, using normalized Raman imaging, Liu *et Al*.^23^ reported that the dry mass of cells is affected quite noticeably by external osmotic stress, where cells grown in hypotonic solutions demonstrated considerably less dry mass than cells grown in isosmotic solutions. This correlates well with the permittivity increase that we detected as a function of decreasing PBS concentration. Consequently, it is reasonable to deduce that the compositional change indicated by the change in ε at microwave frequencies is largely independent of ionic concentration, conductivity, or ionic polarizability, and is instead predominantly dependent on the total polarizability of the intracellular biomaterials relative to water. As the biomaterial-to-water ratio decreases due to water uptake, the permittivity of the cell interior increases. This change in the permittivity signal was further confirmed by analysing cells in a more concentrated (hypertonic) PBS solution. At 150% of the original concentration, we observed a significant decrease in cell geometric size, down by ∼10%, accompanied by a corresponding decrease in relative permittivity of ∼3%. This change in size was due to water expulsion, while the decrease in permittivity reflects the internal biomass-to-water ratio increase due to the loss of water through dehydration.

Conducting single-cell measurements at the microwave band provides a novel mode of probing the internal composition of cells, distinct from impedance cytometry. Here, compositional changes detected by the sensor are analogous to the mass density of a cell, which in this context would be indicated by an ‘electrical density’ value. Given that the biomaterial-to-water ratio, or cell density, is a tightly controlled parameter, significant alterations in the measured ε value can indicate pathological conditions within a cell population or substantial compositional changes due to extraordinary environmental conditions experienced by the cell population.

### Drug resistance probed by Microwave Cytometry

We have established in an earlier work^16^ that the change in cell permittivity at microwave frequencies is a strong indicator of compositional changes inside a cell following the application of a fixation agent. Compositional alterations are commonly observed in cancer cells as well, such as nuclear enlargement leading to an increase in nuclear-to-cytoplasm ratio, as well as hyperchromasia which increases nuclear density.^28,29^ Although solid tumors are stiff, individual cancer cells are softer than non-tumor cells, due to re-arrangement of actin/cytoskeleton networks to allow for flexibility to metastasize.^30^ Such changes in the cytoskeleton are expected to shift the electrical polarizability of cells.

The intracellular structure of cancer cells also changes in response to the presence of physiological stressors such as drugs. In fact, Kimmerling *et al*. were able to show a clear link between drug exposure and changes in the distribution of cellular mass using a SMR mass sensor.^14^ The changes in cellular mass were observed after exposure to over 60 different types of drugs spanning various mechanisms of action over twelve different cancer types. Instead of inferring compositional change inside the cell using a single parameter (buoyant mass), our sensor provides multi-dimensional information about the cell: electrical size (an analogue of dry mass), geometric size (obtained electronically using the same sensor), and microwave permittivity. This capability allows us to monitor internal changes in the cellular structure and content accurately to determine whether a specific cancer cell line is resistant to a specific drug or not. We used isogenic pairs of parental and resistant cell line models to ensure that the difference between sensitive and resistant cells are only due to the drug being investigated. This is critical for our approach as we need to test our platform’s capacity to detect sensitivity and resistance with high precision and dynamic range upon drug treatment.

Here, in the first set of experiments, we used colorectal cancer DLD-1 cell line as the parental (or sensitive) for a series of drug testing experiments. Starting with parental DLD-1 cells (DLD1.Par), we generated isogenic cell line DLD1.GefR which acquired resistance to gefitinib *in vitro* through prolonged exposure to the drug (Methods and Supplementary Information sections).^31–33^

Following the acquisition of gefitinib-resistant populations, both parental and gefitinib-resistant populations were incubated in growth media spiked with increasing concentrations of gefitinib (0, 10, 20, 30, 40, and 50 µM) for 72 h. Figure 3 describes the measurement results of the cell populations for both geometric size and electric size as percentage changes from nominal values for cells in ideal growth conditions using our sensor. Cell viability was measured in parallel (Methods section). The effects of the drug on cell populations can be followed in a two-dimensional plane for geometric and electric size change for cells (Figure 3a). In this representation, the changes in cellular volume such as swelling or shrinkage appear as changes along the x-axis, while the compositional changes in the electrical domain have a projection along the y-axis where higher values represent a higher intracellular biomaterial content and vice versa. For parental (gefitinib-sensitive) cells, even at the 10 µM concentration, we observe a significant swelling in cell geometric size, accompanied by a slight increase in electrical size (Figure 3a, circle markers). This might have resulted from increased water uptake as an adaptation to the drug by the cells. Even at these small drug concentrations (10 and 20 µM), the cell viability (seen as colormap-coded symbols in Figure 3a) appeared to decrease dramatically (∼50%).

**Figure 3:**
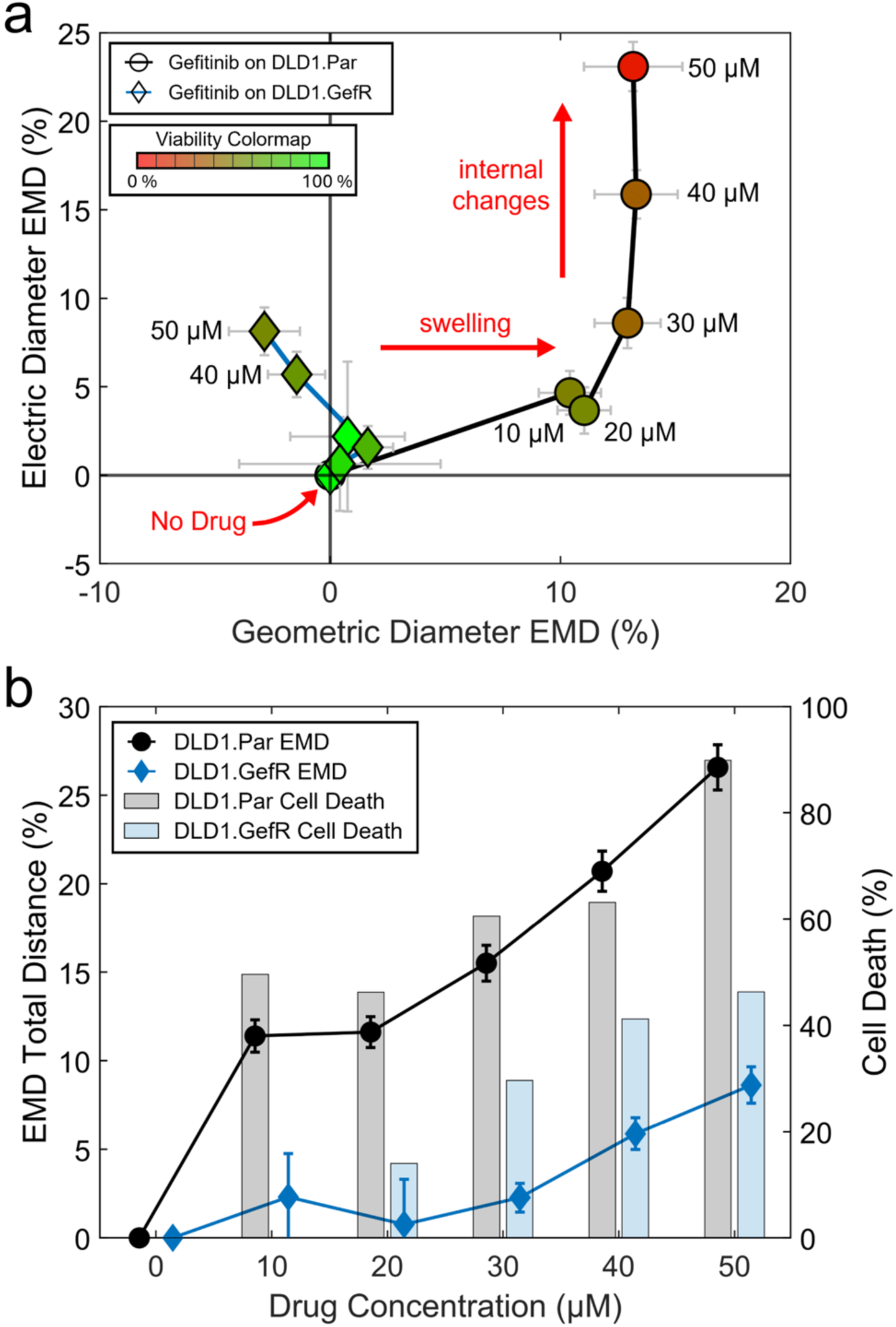
Gefitinib vs parental (DLD1.Par) and Gefitinib-resistant (DLD1.GefR) cell lines. (a) The changes in the geometric diameter (x-axis) and electrical diameter (y-axis) using Earth Mover’s Distance (EMD), for the parental (DLD-1.Par, circle symbol) and gefitinib-resistant (DLD1.GefR, diamond symbol) cell-lines. The drug concentrations vary from 0 µM (No Drug) to 50 µM in a linear fashion as shown. For DLD1.Par cell line, low concentrations of drug predominantly induce a volumetric change, whereas high concentrations cause structural changes as picked up by the electrical diameter. In contrast, the effect of gefitinib on DLD1.GefR cells does not appear to be significant, as expected. Indeed, the induced changes are smaller than statistical uncertainty so long as the concentration is equal to or below 30 µM which is the level at which cells were maintained. The colours in figure markers indicate the cell viability values where bright green indicating 100% and dark red 0% as detailed in colormap legend. Error bars indicate 2 standard deviation levels. (b) Distance to the origin in the EMD plane for the two cell types as a function of drug concentrations (left y-axis), and the cell death percentage (i.e. 100 – cell viability). For each cell type, the total EMD distance runs parallel to the observed cell death levels.

The viability continued to decrease with increasing gefitinib concentrations, reaching 10% at the highest concentration of drug (50 µM). Concurrently, at concentrations >20 µM, the cells maintained their swelling, but the electrical size increased rapidly as well, indicating an increase in cell density. This might be a consequence of cellular hypertrophy.^34–37^ By contrast, this trend was not observed in gefitinib-resistant cells (Figure 3a, diamond markers). In fact, both geometric size and permittivity remained nearly unchanged until concentration values exceeded 30 µM. For concentrations over the 30 µM threshold, a slight decrease in size was observed along with a slight increase in electrical size, which might indicate a loss of water content when coupled with the decrease in size. This is a significant observation, since gefitinib-resistant cells were kept in 30 µM gefitinib media prior to performing the experiment to maintain their tolerance to the drug, so it would be reasonable to assume that the cells will only begin to experience negative effects at concentrations greater than 30 µM. This is further supported by viability measurements where we observe an increase in cell death (defined as 100% – cell viability) from approximately 5% to 20% transitioning from 30 to 40 µM concentration (Figure 3b). Generally, it seems that cell populations that experience more negative effects from drugs tend to exhibit more severe structural changes either in geometric size or in density (electrical size). Therefore, it is useful to look at the extent of the overall change in both dimensions as an indicator of drug susceptibility. Overlaid with the viability measurements in Figure 3b, we also plot the total EMD distance (norm-2 distance of electric and geometric EMD values). The total sensor response and the cell viability measurements run in parallel to each other in this case (Figure 3b) indicating the connection between the electronic measurements and the viability of the cells.

### Cisplatin Response in Gefitinib-Resistant Cells

The results above showed that our sensor system can predict the response of gefitinib-resistant and parental cells under gefitinib treatment by tracking the structural change they experience relative to nominal values in the control samples. Next, we investigated whether a different drug (cisplatin) would have a noticeable effect on gefitinib-resistant cells. Given that the gefitinib-resistant cell line was not necessarily resistant to other drug classes, the exposure to cisplatin (a cytotoxic DNA-damaging drug) is expected to yield dramatic changes in the geometric and electrical size of gefitinib-resistant cell line.

We first exposed the gefitinib-resistant cell line (DLD1.GefR) to 0, 10, 20, 30, 40, and 50 µM of cisplatin as described in the Methods section. We compared the sensor response for the DLD1.GefR cell line with respect to the application of gefitinib *vs.* cisplatin in Figure 4a. In this case, DLD1.GefR cells demonstrated sensitivity towards cisplatin where at the smallest dose (10 µM) as observed by cell death. Indeed, at this dose, we observed a marked increase in the sensor reading (EMD Total Response) compared to the case with gefitinib. As the concentration is increased, the cisplatin-treated cells keep producing larger response in the sensor compared to their gefitinib-treated counterparts, and in parallel with the cell viability measurements superimposed on the graph.

**Figure 4:**
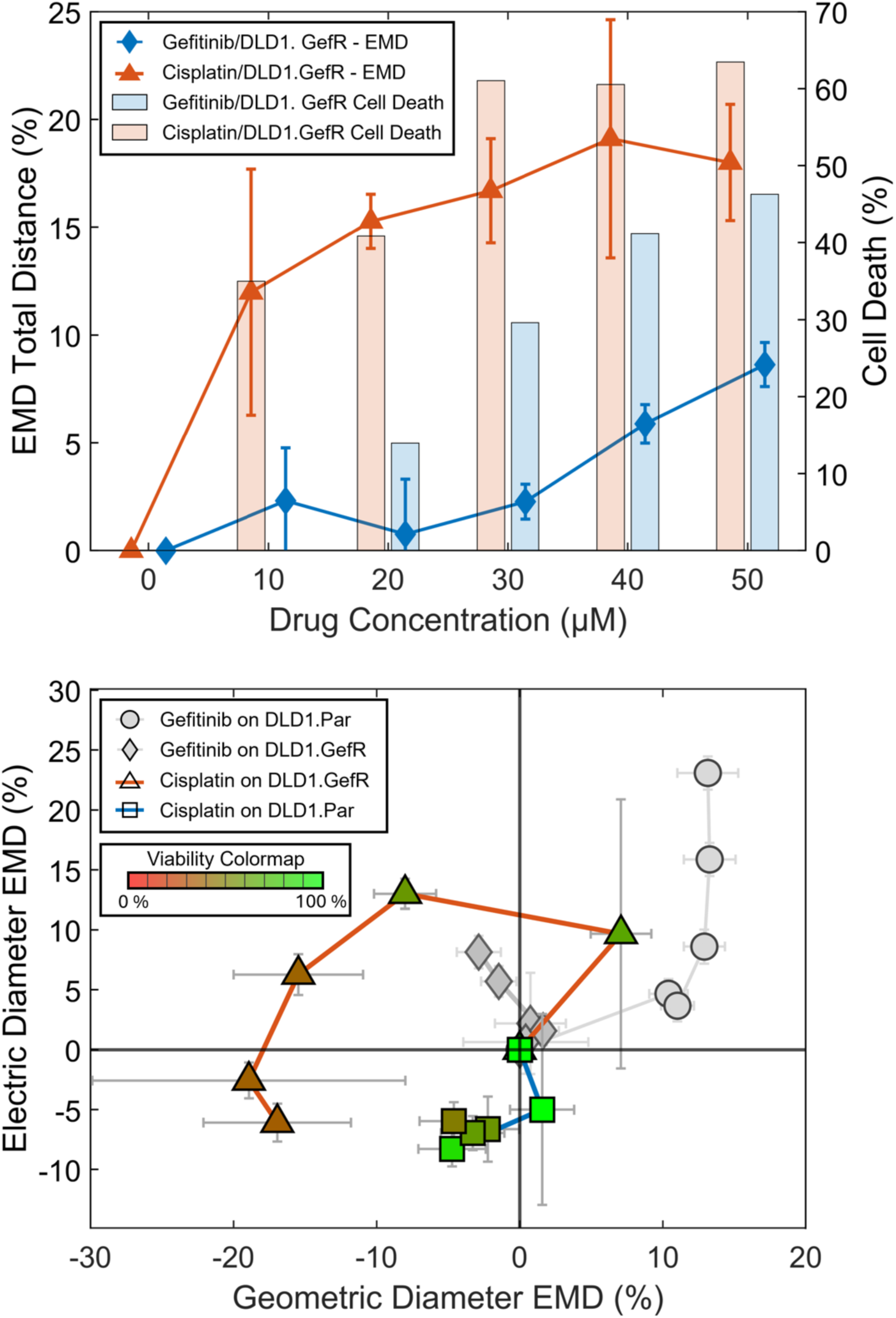
The effect of cisplatin on gefitinib-resistant cell line (DLD1.GefR). (a) Comparison of the response of gefitinib-resistant cell line against two different drugs: gefitinib (blue) and cisplatin (red). When cisplatin is used, both the EMD response and the cell death values attain large values even at low concentrations; on the other hand, the response for gefitinib is smaller for both parameters, especially until 30 µM concentration of gefitinib. (b) The two-dimensional representation of changes in the cell lines as a function of drug. The gefitinib-resistant cell line (DLD1.GefR, triangles) responds to the drug by large shifts in geometric and electrical diameter. The effect of cisplatin on the parental DLD1 cell line (DLD1.Par, squares) is less pronounced, as discussed in the text. The drug concentration levels range from no drug at the origin to 50 µM at the end of the sequence, with 10 µM increments at each step. The colours in figure markers indicate the cell viability values where bright green indicating 100% and dark red 0% as detailed in colormap legend. Error bars indicate 2 standard deviation levels. For ease of comparison, the figure also contains gefitinib treatments discussed earlier in Fig. 1a as grey symbols (circles: gefitinib treatment on DLD1.Par, diamonds: gefitinib treatment on DLD1.GefR).

Figure 4b compares the response of DLD1.GefR *vs.* DLD1.Par under the effect of cisplatin. In this figure, the electrical and geometrical responses of DLD1.GefR *vs.* DLD1.Par cell lines are plotted as a function of cisplatin concentration, superimposed on the results from the previous two experiments (with gefitinib) for comparison. For the gefitinib-resistant cell line (triangles, DLD1.GefR), we see an increase in geometric size (swelling) and an increase in electric volume (the analogue of dry mass). This was followed by a sharp decrease in geometric size for intermediate drug concentrations (20-30 µM) with a small decrease in electrical size, which could be consistent with an apoptotic trajectory where a cell loses between 20-40% of its original volume,^38^ followed by membrane blebbing and the expulsion of intracellular components.^39^ Finally, the changes in cellular volume and electrical size saturate at the highest dose levels where both the size and density of cells are lower than nominal cell values in the control sample. Remarkably, the use of a cytotoxic drug (cisplatin) and cytostatic drug (gefitinib) charts different trajectories in this plane: for instance, cisplatin causes the cells to die as they shrink consistent with an apoptotic trajectory. On the other hand, gefitinib-treated cells die at larger volumes, indicating the result of cell cycle arrest and hypertrophy consistent with the cytostatic nature of the gefitinib.

Parental DLD-1 cell lines (DLD1.Par) exhibited a smaller response, compared to the gefitinib-resistant cell lines (Figure 4b blue trace, square symbols) indicating better tolerance to cisplatin. The tolerance of DLD1.1Par can be understood in the light of a previous study by Fernandez *et al*,^40^ which showed that DLD1.Par cells moderately tolerated ∼40 µM of cisplatin exposure, with 35% of cells remaining viable after 96 h of treatment. Using the aforementioned study as a benchmark, our viability results (Figure SI S2) showed that parental DLD1.Par cells exhibited a 62.9% viability following incubation with 40 µM cisplatin, and at 50 µM cisplatin concentration the viability dropped to 48% after 72 h of drug exposure. This indicates that the DLD1.Par cells tolerated cisplatin at the concentration range applied here. Interestingly, both electrical and geometric size hardly changed across the same concentration range (Figure 4b, square markers). The seemingly unexpected observation where the gefitinib-resistant cell line was more sensitive to cisplatin than the parental cell line could be explained by the possibility of a synergistic effect between gefitinib and cisplatin as detailed in SI S2.

Based on the results so far, we were able to conclude that DLD-1 parental cells would undergo drastic compositional changes both geometrically and electrically when exposed to a drug they were sensitive to. This trend is in line with similar but diminished changes observed in the resistant lines.

### Broad Applicability Across Drugs with Diverse Mechanism of Action

Since cells differ significantly in their response to drugs, we next investigated whether these compositional changes were a universal trait when cells are exposed to drugs that they were sensitive to. To this end, we complemented our earlier results on a cytostatic drug (gefitinib) with cytotoxic (cisplatin) and mixed-effect (tamoxifen) drugs, and corresponding isogenic cell lines. For cisplatin we used the parental HCC-1937.Par and its cisplatin-resistant variant HCC-1937.cisR. For tamoxifen, we used the MCF-7.Par parental cell line and its tamoxifen-resistant variant MCF-7.tamR. The concentration range of each drug was adjusted based on the IC50 concentrations determined for the parental cell lines against each drug (shown in SI Section 5). Given our observation that both geometric and electric sizes exhibited significant changes under the stress of drugs, we decided to quantify the cumulative shifts in both dimensions using the Hotelling T^2^ test which is a measure of the difference between two statistical distributions in two dimensions. Hotelling T^2^ scores provide a single value that is commensurate with changes in both of those dimensions relative to the origin point (control sample). The results for the Hotelling T^2^ for each drug/pair combination are shown in Figure 5 including the previously tested DLD1 isogenic cell lines to allow comparison. In Fig. 5, a large value of Hotelling T^2^ (bar plots) indicates a large change in the electrical and geometric diameters of the cell line under the influence of the applied dose of the drug. As visible from Figure 5, in all cases, there is a large Hotelling T^2^ peak for the parental (drug sensitive) cell lines which indicates a sudden change in biophysical parameters at the applied drug level. Indeed, the Hotelling T^2^ peaks emerge near concentration values where cellular viability has a sharp drop (which in turn are often close to the LC_50_ values). These results on different cancer types further indicate the utility of our technique for predicting the response of the drug.

**Figure 5:**
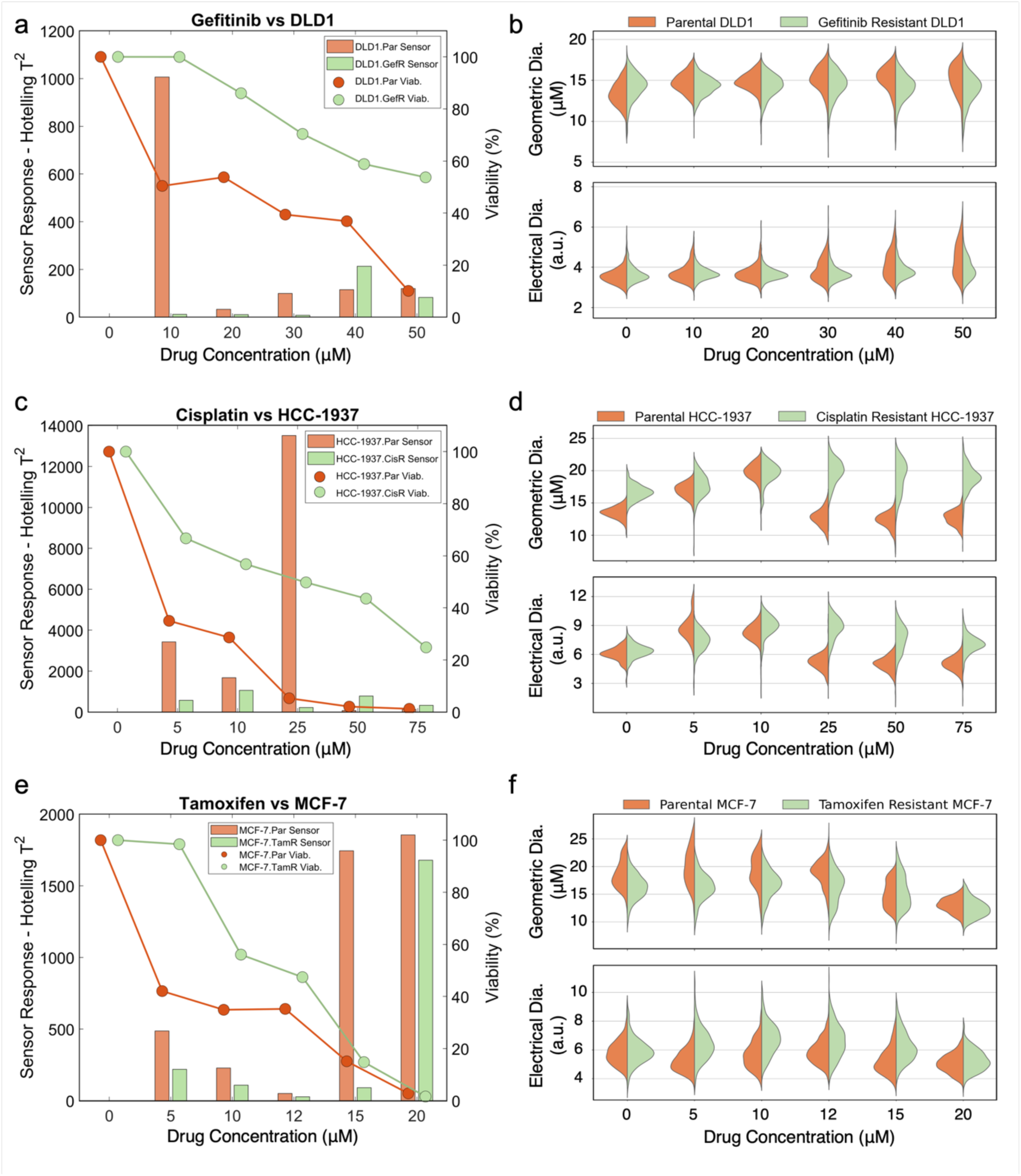
Left Panels: Hotelling T^2^ statistical test results (left-axis, bar plots) and viability values (right-axis, dotted-lines) for three different drugs and their corresponding parental (sensitive) and resistant cell lines, in orange and green respectively. Right Panels: Electric and geometric diameter distributions obtained by our sensor as a function of applied drug concentration. The tested drugs and cell lines are a),b) Gefitinib on DLD1, c),d) Cisplatin on HCC-1S37, and e),f) Tamoxifen on MCF-7. The larger Hotelling values indicate a large shift in the biophysical parameters in the cells. As evident from the figure, largest shifts in Hotelling occurs in the sensitive parental cell lines near the dose levels where there is a large drop in viability. The Hotelling T^2^ values are calculated sequentially, i.e. each population is compared to the population at the previous (smaller) drug level. The Hotelling T^2^ values in part (c) and(e) have initial large peaks close to their LC_50_ value, and larger peaks later on at drug concentrations where the cell viability almost goes to zero.

Examining population distributions in both geometric and electrical dimensions offers detailed insights into the evolution of cell populations under drug influence. This is visualized in the violin plots for each drug/cell pair (Fig. 5b, d, and f). Key indicators include abrupt shifts in population distribution peaks and changes in population dispersity or broadening.

In the gefitinib vs. DLD-1 violin plot (Fig. 5b), we observed no significant changes in geometric or electrical diameters, except for the parental cell population within the 0– 10 µM range, which corresponds to a pronounced Hotelling peak. Another notable feature is the widening of the electrical diameter peak for the parental population at 30 µM, mirrored by a similar widening in the resistant population later on at 50 µM.

Given gefitinib’s cytostatic nature, we anticipate more pronounced compositional changes in cell populations exposed to a cytotoxic drug like cisplatin. Indeed, the violin plots for DLD-1.GefR vs cisplatin (SI, Section 3) do show drastic geometric shifts and extensive peak broadening starting at the 10 µM concentration. These shifts are much more drastic than DLD-1.Par vs gefitinib. We also see peak broadening in the electrical domain for DLD-1.GefR vs cisplatin starting from the lowest concentration, compared to minimal peak broadening in the electrical domain for DLD-1.Par vs gefitinib except at concentrations of 30 µM and larger.

Similarly, the application of the cisplatin on HCC-1937 indicates dramatic reductions in both geometric and electrical diameters are evident for parental cells at 25 µM drug concentration (violin plot in Fig. 5d). These drops are preceded by gradual increases in size at lower drug concentrations, with the highest Hotelling peak occurring at 25 µM for the parental line (Fig. 5c). In contrast, resistant cells exhibit no such sharp decreases; however, significant increases in dispersity suggest a heterogeneous response within the resistant population, where some cells experience adverse effects while others tolerate the drug better.

For the application of tamoxifen on MCF-7 cells, we see that at low dose levels (5 µM and 10 µM), the MCF-7.Par cell line undergoes a quick loss of viability, dropping below 40% (Fig. 5e). Within this range, we see that the viability of the MCF-7.TamR stays at relatively higher levels. These observations are recapitulated in the sensor response curves, as the MCF-7.Par response value is almost double, compared that of MCF-7.TamR. At the highest concentration levels (>15 µM), viabilities converge to 0 % for both types (i.e. all cells die); as such, these dramatic drops in both types exhibit very large sensor responses. This is reminiscent of the earlier case with HCC-1937.Par response at 25 µM concentration of cisplatin (Fig. 5c), where the full extinction of cells was accompanied by a large change in sensor response. The geometric and electrical diameter distributions for the MCF-7.Par and MCF-7.TamR cell lines are provided in Fig. 5f.

The distribution patterns of electrical and geometric diameter reveal a key distinction: resistant cells tend to maintain the overall geometric and electrical size under drug exposure except when the drug dose is too high, while parental cells display rapid deformations, such as sudden changes in geometric and electrical size, coupled with distribution broadening or splitting into sub-populations. Notably, resistant cells also exhibit distribution broadening and population splitting especially at higher dose levels, which might indicate a heterogeneous response to the drug among the resistant population, especially at higher drug doses.

These findings provide valuable insights into the mechanistic effects of specific drugs on various cell lines, enabling a deeper understanding of drug susceptibility and resistance in terms of their effect on cellular composition.

### Measurements on Patient-Derived Organoids

To further generalize our observations, we applied our technique on patient-derived organoids (PDOs), highlighting its potential for translational applications.^41^ PDOs provide a model system that closely mimics the heterogeneity of tumours.^42^ By using cells directly from patients, this approach demonstrates an accurate investigation of their therapeutic response.^43^ Generally, primary cells grown in 3D organoid cultures are expected to recapitulate the response *in vivo* more faithfully than conventional 2D cell cultures owing to their growth conditions and geometry.^44^ As a result, the demonstration of microwave cytometry on PDOs serves as an important proof-of-concept towards clinical applicability.

The effects of Gefitinib and Bortezomib on the cell viability of the PDOs from the same donor were previously assessed using the Sytox-Green nucleic acid staining procedure, followed by 3D nuclear visualization for quantitative analysis (manuscript in preparation). Treatment with 1 µM Gefitinib and 1 µM Bortezomib resulted in cell viabilities of ∼100% and <10%, respectively, compared to the control group treated with DMSO. Consequently, three organoid groups from the same donor were grown in optimal conditions (Methods) and two of those groups were then incubated with gefitinib and bortezomib respectively at concentrations of 1 µM, while the third was maintained in optimal conditions and served as control. Following the growth and incubation periods, the organoids were dissociated into single cells through gentle digestion using TrypLE^TM^ and then passed through the sensor.

The analysis revealed distinct differences in the cell populations. While the control population and the gefitinib-treated population aligned closely along the 2D geometric versus electric diameters plot (Fig. 6a), cells treated with bortezomib were noticeably smaller in size. A clear trend emerged: gefitinib-treated cells were slightly smaller in both geometric and electric diameters compared to the control, while bortezomib-treated cells were significantly smaller in both parameters.

**Figure 6:**
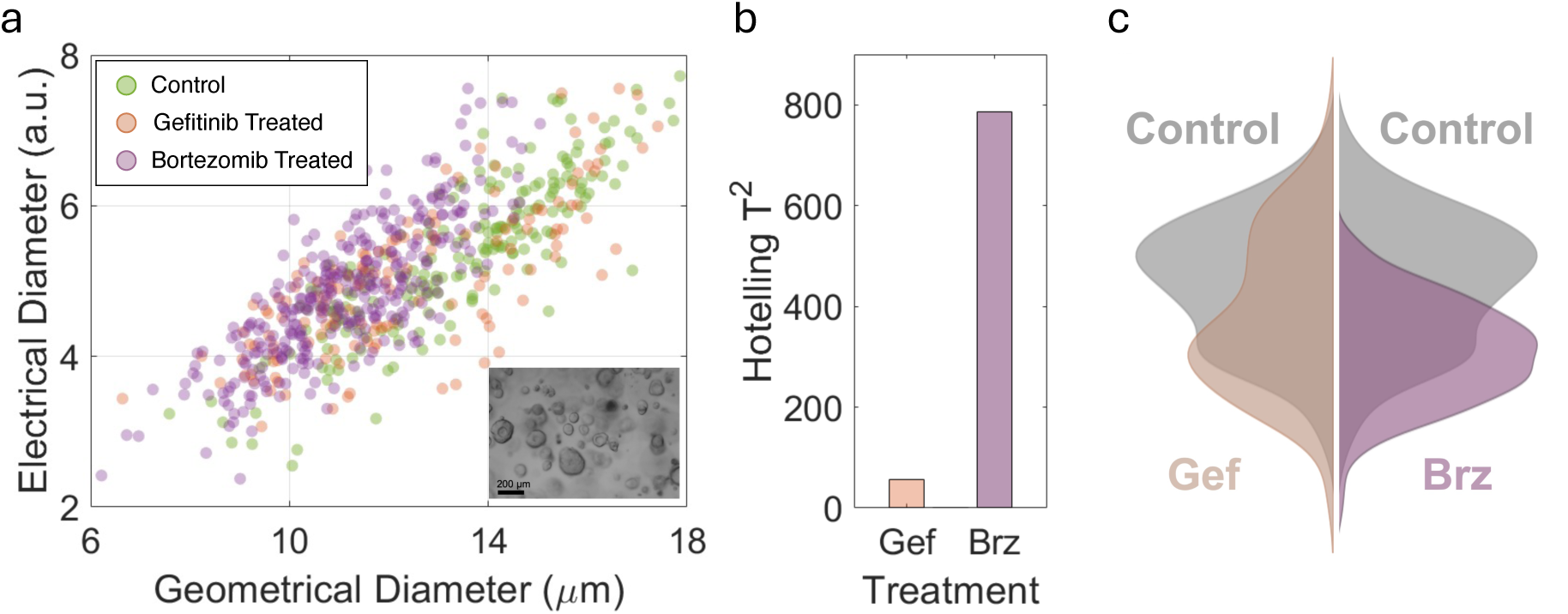
Measurements on cells isolated from PDOs. (a) The electrical vs geometrical diameter of cells. The gefitinib treated cells (blue) overlap mostly with the control group (green), while the bortezomib treated cells have smaller geometric diameters and slightly larger electrical diameters. Inset shows a general picture of organoids used in the experiments. (b) The Hotelling T^2^ test clearly indicates that the changes in the bortezomib treated group are much larger. Inset shows the geometric diameter distribution for the control (grey), gefitinib-treated (orange) and bortezomib-treated (purple) groups. (c) The geometric diameter distribution for the control (grey), gefitinib-treated (orange), and bortezomib-treated groups.

Statistical analysis using the Hotelling T² test confirmed significant variation in both geometric and electric diameters for the bortezomib-treated population compared to the control (Fig 6b). In contrast, the gefitinib-treated population showed only minor variations relative to the control group. Observing the data at finer granularity, the violin plot (Fig 6c) shows that the gefitinib-treated sample shows two sub populations where possibly some individual cells tolerated the drug better than their counterparts in the other sub population. This becomes clearer when compared to the bortezomib-treated population where a single population appeared with a peak value that deviated sharply from the control sample, indicating that almost all the cells in the population experienced adverse effects due to their exposure to the drug.

## Conclusions

In this work, we have shown for the first time that the combination of microwave and low-frequency electronic measurements could be used to determine the susceptibility of a cell line to a given drug. First, we validated the sensor’s ability to measure changes in the biomaterial-to-water content of cells by exposing them to salt solutions with different osmolarities. We then conducted our drug susceptibility study using a pair of DLD-1 cell lines — one of which was sensitive and the other resistant to gefitinib. The effects of gefitinib on these cell lines were successfully monitored across multiple biophysical dimensions. The parental cell line exhibited swelling and significant alterations in microwave permittivity which is indicative of changing biomaterial-to-water ratio. In contrast, the resistant cells showed almost no change in sensor response, even at concentration up to 30 µM —the level at which they were incubated to maintain drug resistance. By contrast, administering a different drug, cisplatin, to the gefitinib-resistant cell line had an immediate effect in the sensor response.

Subsequently, we applied this methodology to additional isogenic cell lines and associated drugs to demonstrate applicability in the major drug mechanisms: cytostatic (gefitinib), cytotoxic (cisplatin) and mixed-effect (tamoxifen) drugs. In each case that the electronic measurements could distinguish between the sensitive and resistant cells. In all cases, the response of the sensor was validated through cell viability measurements. Moreover, the ability to disentangle volumetric and compositional changes, and the analysis of the effects of cytostatic and cytotoxic drugs on cells tested here, indicate the potential of microwave cytometry for identifying mechanisms of actions for different drugs.

Finally, we showed the potential for translational research and clinical application by analysing cells from PDOs. Our findings demonstrate that this approach can provide important insights into cell viability of primary cells in PDOs better reflecting tumour heterogeneity, offering a potential clinical application. Other biological samples, such as blood, pleural effusion and fine-needle aspirates can also be used in our system after isolation of cancer cells but without the need for culturing, since our microfluidics platform and single-cell technology can provide drug-resistance information by using only several thousand cells.

Given the versatility of our system in testing diverse drugs on single cells from both cultured cells and PDOs, this approach offers a powerful avenue to tailor patient-specific treatment modalities, advancing precision medicine and ultimately improving clinical outcomes.

## Materials and Methods

### Sensor Device Fabrication

The sensor device was comprised of a planar split-ring resonator (SRR) design deposited on fused silica wafers (Quartz Unlimited LLC). Device fabrication has been described in detail in previous publications.^16,45^ Briefly, standard soft lithography was utilized to deposit a thin Au layer to form the body of the SRR using a pre-etched Cr photolithography mask. The SRR comprised of two concentric rings, one larger outer ring, and another smaller inner ring. The outer ring was split, with two Au traces coming out of each end and terminating with a sub-miniature A (SMA) connector soldered onto the edge of the fused-silica wafer. The inner ring comprised of another split circle where the split comprised of a small region where signal sensing took place. The sensing region in the inner ring consisted of two asymmetrical gaps spanning a length of approximately 250 µm across (Figure S1). Furthermore, two Au electrodes were deposited surrounding the SRR sensing region and extending outwards for subsequent wire bonding. These electrodes were connected to a low frequency signal generator and acted as a conventional Coulter counter. Finally, a straight (100 µm wide, 45 µm tall, and 70 mm long) microfluidic channel with a constriction (40 µm wide, 45 µm tall, 250 µm long) in the centre was developed in polydimethylsiloxane (PDMS, Dow Chemicals) featuring an inlet and outlet at either ends. This channel was then carefully aligned and bonded (through O_2_ plasma treatment) so that the constriction in the channel covered the sensing region of the SRR precisely.

Liquid containing samples was delivered through the microfluidic channel using a Fluigent (MFCS-EZ) pressure control system. Typical flow rates ranged from 8 to 15 µL/min with a cell transit time through the SRR sensing region being between 10-20 ms.

### Electronic Measurements

The sensor platform consists of a low-frequency and a high-frequency (microwave) sensor to obtain the internal property of the cells. In the low-frequency part, referred as Coulter counter, an insulating particle’s passage results in partial blockage of ionic current being conducted between two electrodes, the decrease in current being proportional to the geometrical volume of the particle. A 0.5 V_pp_ signal at 0.5 MHz from a lock-in amplifier (Zurich Instruments, HF2LI) was used to drive one electrode while the resulting current flowing through the ionic solution was collected from the other electrode. This signal was converted to voltage by a transimpedance amplifier (Zurich Instruments, HF2TA) and consequently read out by the lock-in amplifier.

The high-frequency part of the sensor utilized a split ring resonator (SRR) for electrical size measurement of the cells. The SRR contains two concentric ings, the inner ring is excited inductively by the microwave signal fed through the outer ring. A highly concentrated electric field is created in the gap (split region) of the inner ring due to the standing wave mode^46–48^ and passage of particle through the gap cause phase and amplitude change of the resonator due to capacitance change in the sensing region representative of the particle’s geometric volume and the permittivity. In this specific design, time delay of the signals from two gaps of unequal width (15 and 25 µm) in the inner ring were utilized to obtain the height information of the passing particles and for calibrating the signal *(Figure S1)*.

A vector network analyzer (VNA) was used to observe resonance characteristics of the SRR before connecting to the custom measurement circuitry for conducting measurements.^16^ A signal generator was used as a signal source and its frequency was set as the resonance frequency of the SRR (≈5.3 GHz). A 0.5 V_pp_ signal at the resonance frequency was fed to the SRR. Two lock-in amplifiers (Zurich Instruments, MFLI) were used to employ single side band modulation (SSBM) as part of the measurement circuitry. A custom-built single side band (SSB) heterodyne circuitry^49–51^ was used as the resonance frequency was higher than the maximum operational frequency of the lock-in amplifiers. Up-conversion and down-conversion were utilized for digitally reading the phase and amplitude of the signal. To obtain signals from fast moving cells, the time constant of the lock-in amplifiers was set as 501 µs, and the sampling rates were set as 14.39 kSa/s and 13.39 KSa/s for low-frequency sensing, and high-frequency sensing, respectively.

Signal obtained from the low-frequency part of the sensor provided the cell’s geometrical volume upon height calibration by the use of calibration particles (Polystyrene 20 µm, Sigma-Aldrich) added into the sample for this purpose. The phase and amplitude response of the microwave resonator were obtained from the high-frequency part of the sensor, and the out-of-phase component (Y ≡ R sinθ) of the reflected voltage, was calculated which probes any minute change in the capacitance of the resonator.

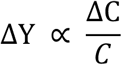

Here, ΔY refers to the change in the out-of-phase component of the reflected voltage, C refers to the total capacitance of the resonator, and ΔC refers to the change of capacitance due to the passage of a cell. The change of phase (Δθ) and amplitude (ΔR) were extracted for transit of single cells through the sensing area. The capacitance change due to the presence of a cell can be calculated by

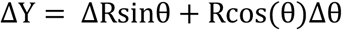

where R and θ refer to the baseline amplitude and phase values in raw signal. The capacitance change calculated is a function of geometrical volume and the

Clausius-Mosotti factor of the cell (a factor depending on the microwave permittivity values of the cell and the medium), decoupling the geometrical volume as obtained by the Coulter counter enables attaining the microwave permittivity of individual cells. The details of the circuitry, calibration procedure, and data analysis can be found in an earlier work.^16^ The Earth-Mover Distance calculations follow the standard definition^14,25^ and the error bars are calculated using bootstrapping techniques using 200 resamples.

#### Cell Culture and Reagents

Human colorectal cancer cells (DLD1) were gifted by Prof. Alain Chariot, GIGA center, Belgium and cultured according to supplier’s protocol. HEK293 FT cells were cultured in Dulbecco’s Modified Eagles’ Medium (DMEM, Biowest), while DLD1 cells were cultured in RPMI 1630 media (Biowest). Subculturing and passaging were carried out using 1x Trypsin EDTA (Biowest) and 1x PBS (Biowest). Parental and cisplatin resistant HCC1937 cells were cultured in DMEM with 10% FBS, 50 U/mL penicillin/streptomycin, 1% nonessential amino acids (Gibco). Parental and tamoxifen resistant MCF-7 cells were cultured in phenol red–free DMEM (Gibco) with 10% FBS, 0.1% insulin, 50 U/mL penicillin/streptomycin, 1% nonessential amino acids (Gibco).

#### Hyper/Hypotonic PBS Microwave Measurements

Hypotonic solutions were prepared by taking commercial sterile 1x PBS solution (Biowest) and diluting it using HPLC-grade water (Sigma-Aldrich) at 1:1, 1:4, and 1:10 PBS:water to gain solutions with 100% PBS concentration, 50%, 20%, and 10%, respectively. A 150% PBS solution was prepared by dissolving 16.9 mg/mL of pure KCl in stock (1x) PBS.

HEK293 FT were grown from frozen stock in 6-well plates in DMEM supplemented with 10% Fetal Bovine Serum (FBS). The cells were grown under incubation conditions for 48 h until reaching 80% confluency. After reaching confluency, the cells were trypsinized using a 1x trypsin-EDTA solution for around 3 min, followed by gentle mixing and aspiration into 15 mL Falcon tubes. The cells were then centrifuged, the supernatant was removed, and 3 mL of fresh PBS solution of either 150, 100, 50, 20, or 10% (as outlined in the results section) was added. The cells were resuspended in the PBS solution through gentle mixing and aspiration. The cells were then injected into the microfluidic channel of the sensor for measurements.

#### Development of Drug-Resistant Cell Lines

Gefitinib was purchased from Santa Cruz Biotechnology, Inc and dissolved in cell culture-grade dimethyl sulfoxide (DMSO). DLD1 cells were continuously exposed to Gefitinib by gradually increasing the concentration over time. Since gefitinib was delivered in DMSO, control DLD1 cell population was cultured in parallel with equal amounts of DMSO without gefitinib, serving as the control cells. Gefitinib resistant DLD1 cell line was obtained when their LC_50_ to the drug reached double the concentration for that of the control cells. It took roughly one year starting from 12 µM,= then the resistant cells were maintained in 30 µM Gefitinib. Cisplatin was obtained from NOVAGENTEK (Turkey) in powder form and working stock solutions were prepared in HPLC-grade water (Sigma-Aldrich). HCC-1937 cisplatin resistant cells were developed by culturing the cells in the presence of 7.5 µM, for over 6 months. The parental counterparts were cultured in the absence of cisplatin. Tamoxifen-resistant MCF-7 cells (MCF-7 TamR) were generated by culturing the cells in the presence of 5 μmol/L of 4-hydroxytamoxifen (Sigma-Aldrich) for over 1 year. In parallel, parental MCF-7 cells were maintained under identical conditions without tamoxifen.

#### Cell Viability Measurements

In parallel with sensor measurements, conventional cell viability measurements were conducted to validate the sensor data. In order to calculate the total cell death, we first calculated the total number of cells remaining in a given sample after the administration of the drug dose and incubation for 72 hours using manual cell counting on a haemocytometer in the presence of Trypan Blue. Cells that took up Trypan Blue were excluded from the total count since they were considered non-viable. The number of remaining cells in the given sample was then compared to the number of cells in the control sample (where 0 µM of the drug was administered). This yielded a percentage of the number of surviving cells in the measured sample relative to the control sample, which was then reported in the text.

#### Drug Resistance Tests - Electronic Measurement Workflow

For microwave measurements of drug-resistant cells and their control ones, cells were seeded into 6-well plates (Greiner Bio One) to obtain 60% confluency next day and allowed to attach to the plates overnight under incubation conditions (37° C, 5% CO_2_,). The following day, the wells were replaced with fresh media with different drug concentrations as described in the figure legends. The plates were then incubated for 72 hours before initiating measurement procedures.

After 72 hours, each well was treated as follows: media containing the drug was aspirated and placed in a falcon tube. A trypsin solution was then added to the well and allowed to incubate at 37° C for approximately 3 minutes. Upon cell detachment, the cells were resuspended in solution through gentle pipetting, and the cell suspension was then recombined with the previously aspirated media in the falcon tube. This was done to collect both floating and adherent cells in the well upon treatments. The collected cells were then centrifuged at 1500 rcf (Eppendorf centrifuge) and the supernatant was discarded. Finally, each of the cell pellets were resuspended in 3 mL of 1 x PBS. 20 µL of the cell suspension was taken and mixed at a 1:1 ratio with filtered (0.22 µm polycarbonate filters) Trypan Blue dye for 1 min, followed by measuring cell viability as described previously using a haemocytometer. The remaining cell suspension was then mixed with 20 µm monodisperse polystyrene microsphere solution (Sigma-Aldrich, Product No: 74491) to serve as an internal calibration standard. The cell suspension was then introduced into the inlet of the microfluidic channel for the sensor device using a Fluigent pressure control system. The cells and polystyrene spheres were allowed to freely flow through the sensing region of the device and up to 500 individual measurement events were collected for each repetition. Three separate viability measurement repetitions were performed for each well. The outflow was then discarded appropriately.

Prior to the subsequent measurement, the microfluidic channel of the device was thoroughly washed with PBS and HPLC-grade water to prevent cross-contamination between samples.

#### Establishment and Culture of PDOs

Patient-derived organoids (PDOs) were generated from a fresh surgical specimen obtained from a patient who underwent surgical resection at Hacettepe University Hospital. The study was approved by the Ethical Committee of Hacettepe University (reference number: KA-20136). PDO establishment and culture were performed as previously described.^43^

Briefly, fresh tissue was transferred into cold Dulbecco’s Phosphate-Buffered Saline (DPBS; Sartorius) supplemented with 1X Penicillin-Streptomycin (Thermo Fisher Scientific). The tissue was washed at least three times with cold DPBS, minced, and collected into 15 mL Falcon tubes. It was then conditioned with 5 mL PBS containing 5 mM EDTA (Sartorius) for 15 minutes at room temperature. Enzymatic digestion was performed using TrypLE™ (Thermo Fisher Scientific) for 1 hour at 37°C. To facilitate cell release, mechanical dissociation (pipetting) was applied. Dissociated cells were collected in Advanced DMEM/F12 (Thermo Fisher Scientific) and filtered through a 40 µm strainer (Greiner) to remove undigested tissue fragments.

The filtered cells were centrifuged at 1200 rpm for 5 minutes at 4°C, resuspended in 120 µL growth factor-reduced (GFR) Matrigel (Corning), and seeded into a 24-well cell culture plate (Corning). The Matrigel was allowed to solidify for 20 minutes at 37°C in a 5% CO₂ incubator, after which 500 µL of Advanced DMEM/F12 medium containing 1X Penicillin-Streptomycin, 2 mM L-Glutamine (Thermo Fisher Scientific), 100 µg/mL Primocin (Invivogen), 1X B27 supplement (Gibco), 1X N2 supplement (Gibco), 0.01% BSA (Sigma), 50 ng/mL human EGF (Biolegend), 100 ng/mL human Noggin (Biolegend), 500 ng/mL human R-Spondin-1 (Biolegend), 10 nM Gastrin (Sigma Aldrich), 1 µM Prostaglandin E2 (Tocris Bioscience), 10 ng/mL human FGF-10 (Biolegend), 10 ng/mL human FGF-basic (Biolegend), 100 ng/mL human Wnt-3a (RCD Systems), 4 mM Nicotinamide (Sigma Aldrich), 0.5 µM A83-01 (Tocris Bioscience), and 5 µM SB202190 (Sigma Aldrich) was added. Additionally, the final medium was supplemented with 10 µM ROCK inhibitor Y-27632 (Sigma Aldrich).

#### PDO Drug Sensitivity Measurement

For the drug assay, PDOs were mechanically harvested using 5 mL PBS/EDTA (1 mM) containing 1X TrypLE™ (Gibco) and incubated for 10 minutes at 37°C. PDOs were then mechanically dissociated into single cells, washed with Hank’s Balanced Salt Solution (HBSS; Thermo Fisher Scientific), pelleted at 1200 rpm for 5 minutes at 4°C, resuspended in 120 µL GFR Matrigel, and reseeded at a density of 1 × 10⁵ cells per well into a 24-well plate. Following Matrigel solidification for 20 minutes at 37°C in a 5% CO₂ incubator, 500 µL of complete organoid medium was added to each well. The medium was refreshed after 24 hours.

Four days post-seeding, 1 µM Gefitinib (Selleckchem), 1 µM Bortezomib (Selleckchem), or DMSO (control) was added to the wells. After 6 days of drug treatment, PDOs were dissociated into single cells using 1X TrypLE™, pelleted at 1200 rpm for 5 minutes at 4°C, resuspended in HBSS, and introduced into the microfluidic device channels for further analysis.

## Acknowledgment

The authors thank Aykut Koc, Alper Demir and Onur Cizmecioglu for useful discussions.

