## Supplemental Figures for "Single-Cell Microwave Cytometry for Drug Resistance Detection in Cancer"

**Section 1. Sensor Technology**

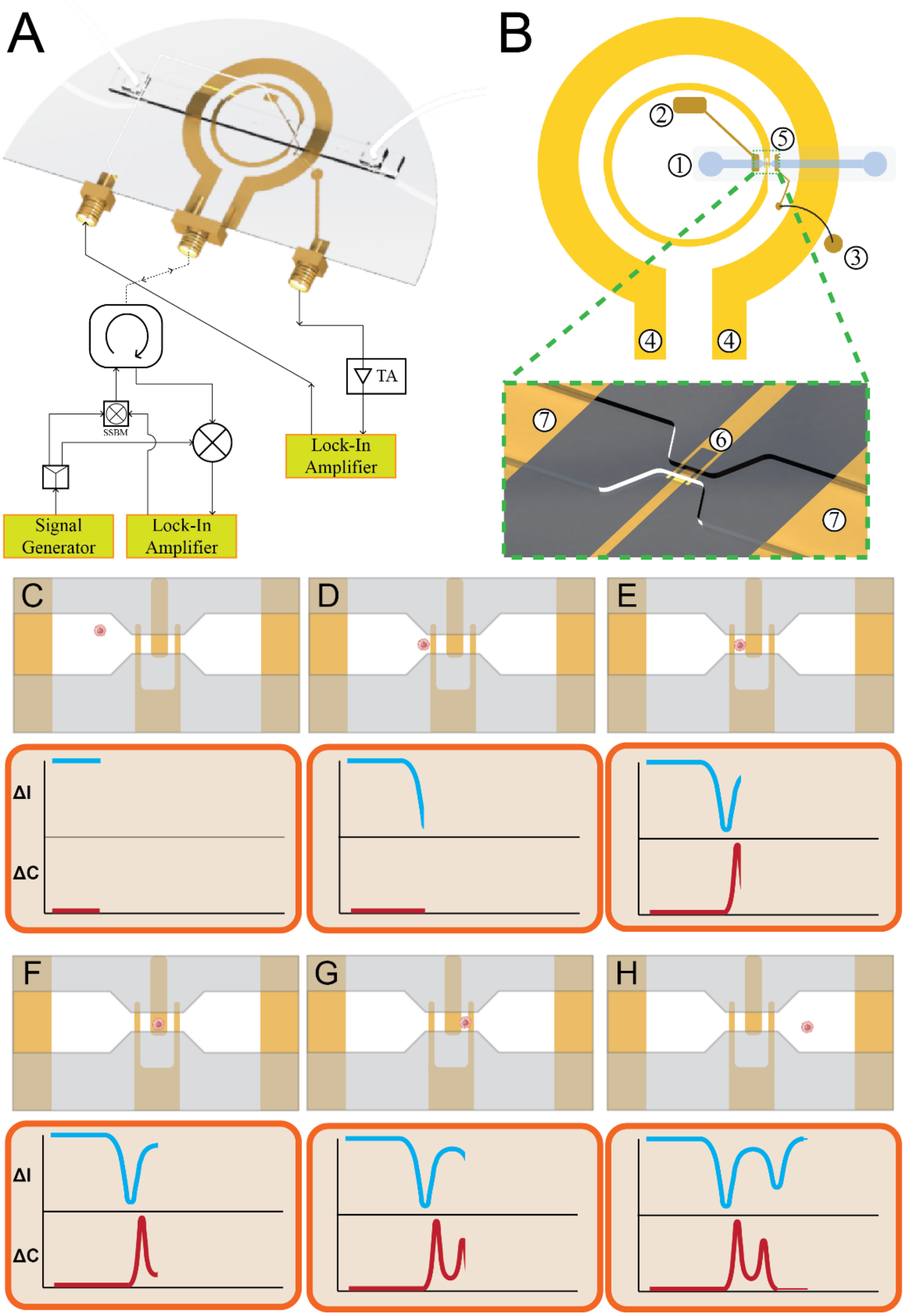


*Figure S1: Schematic diagram representing the sensor architecture and operation where A) is a 3D render of the platform showing the microchannel spanning the inner and outer rings of the SRR, with the gold pads forming the Coulter counter also visible. It also shows a simple diagram of the wiring scheme for each sensor. B) A schematic view of the same SRR/Coulter arrangement where (1) is the channel, (2) is the left electrode of the Coulter counter, (3) is the gold pad that is bonded to the right*

*Coulter counter electrode, (4) is the outer SRR ring , (5) is the sensing region containing the gap in the inner ring, while the inset is a close up of the gap and sensing region clearly showing (6) the inner ring gap aligned with the constriction in the microfluidic channel and (7) the outer electrodes forming the Coulter counter and connected to (2) and (3). C-H) Are a series of schematics showing a cell passing through the sensing region, and how its passage generates a signal response in both the current (ΔI) from the Coulter sensor and the capacitance change (ΔC) from the microwave sensor that represent geometrical, and electrical sizes respectively. As the cell nears the constriction, it blocks the current passing between the Coulter electrodes appearing as a reduction in conductivity. When the cell reaches the split-ring gap, it significantly alters the capacitance of the SRR structure especially that the electric field is concentrated at this point, therefore generating its own signal. Both those signals are then processed to calculate the electrical permittivity of the cell that passed through.*

### Section 2. Resistance of DLD1 Parental Cell Line to Cisplatin

The seemingly unexpected gefitinib-resistant cell line sensitivity to cisplatin (Figure 4b in main text, and S2 below) could be explained by the possibility of a synergistic effect between gefitinib and cisplatin. Previous studies have shown that EGFR-TKIs (such as lapatinib) chemosensitize ovarian cancer cells to cisplatin,[1] while EGFR-inhibitor osimertinib combined with pemetrexed or cisplatin showed effective tumour suppression in lung cancer.[2] Ahsan *et al*. showed that cisplatin induced EGFR phosphorylation and degradation in wild type human head and neck carcinoma.[3] Furthermore, they also showed that this degradation effect was enhanced by the administration of EGF, thus cells that rely on that receptor for growth (which is the case with gefitinib-resistant DLD-1) might be more susceptible to EGFR degradation through cisplatin-induced cytotoxicity. It is also worth noting that a randomized phase III study, concluded in 2017, evaluated pemetrexed/cisplatin as a first-line therapy for patients with non-small cell lung cancer tumours with EGFR-activating mutations (NCT00949650).[3] This leads us to assume that the enhanced sensitivity to cisplatin exhibited by gefitinib-resistant cells might be due to EGFR degradation. In this specific case, both geometric size and permittivity plummeted at the first administered cisplatin dose, and continued to fall further, possibly indicating an apoptotic trajectory for death, highlighted by a rapid loss of size and the condensation of the biomaterial content.

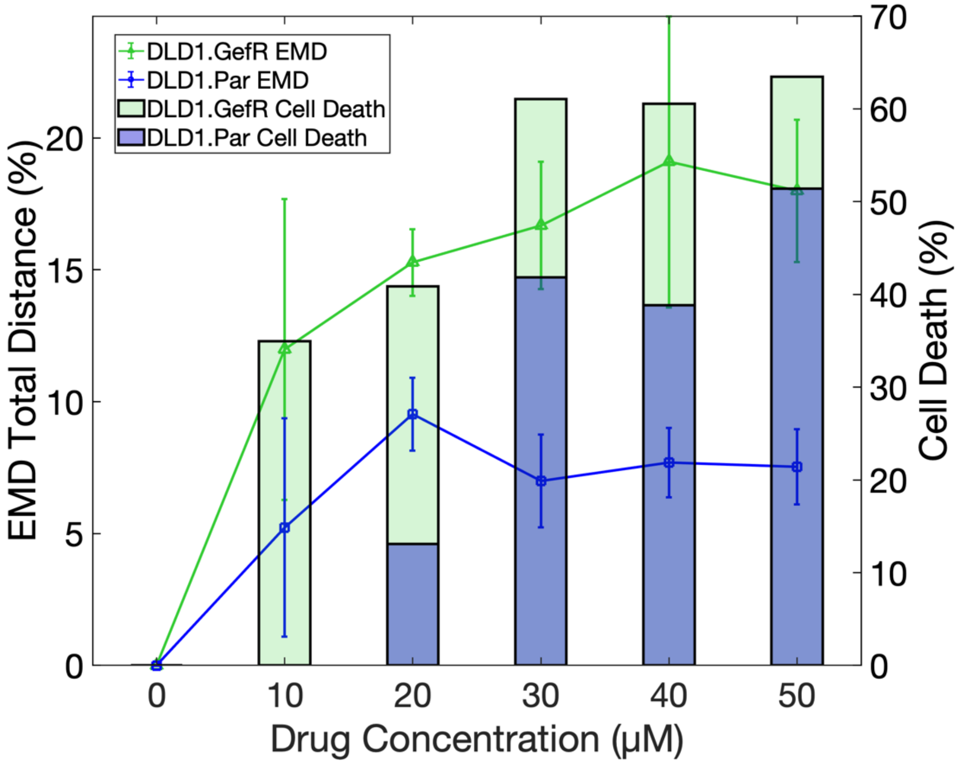


*Figure S2: For cisplatin treatment, comparison of the parental DLD1 cell lines (blue, DLD1.Par) and gefitinib-resistant cell lines (green, DLD1.GefR) in terms of total EMD distance to the origin and cell death. Both parameters indicate that gefitinib-resistant cell line is more sensitive to cisplatin compared to the parental DLD1 cell line, as explained in the test*

### Section 3. Cell Distributions under Drug – Split Violin Graphs

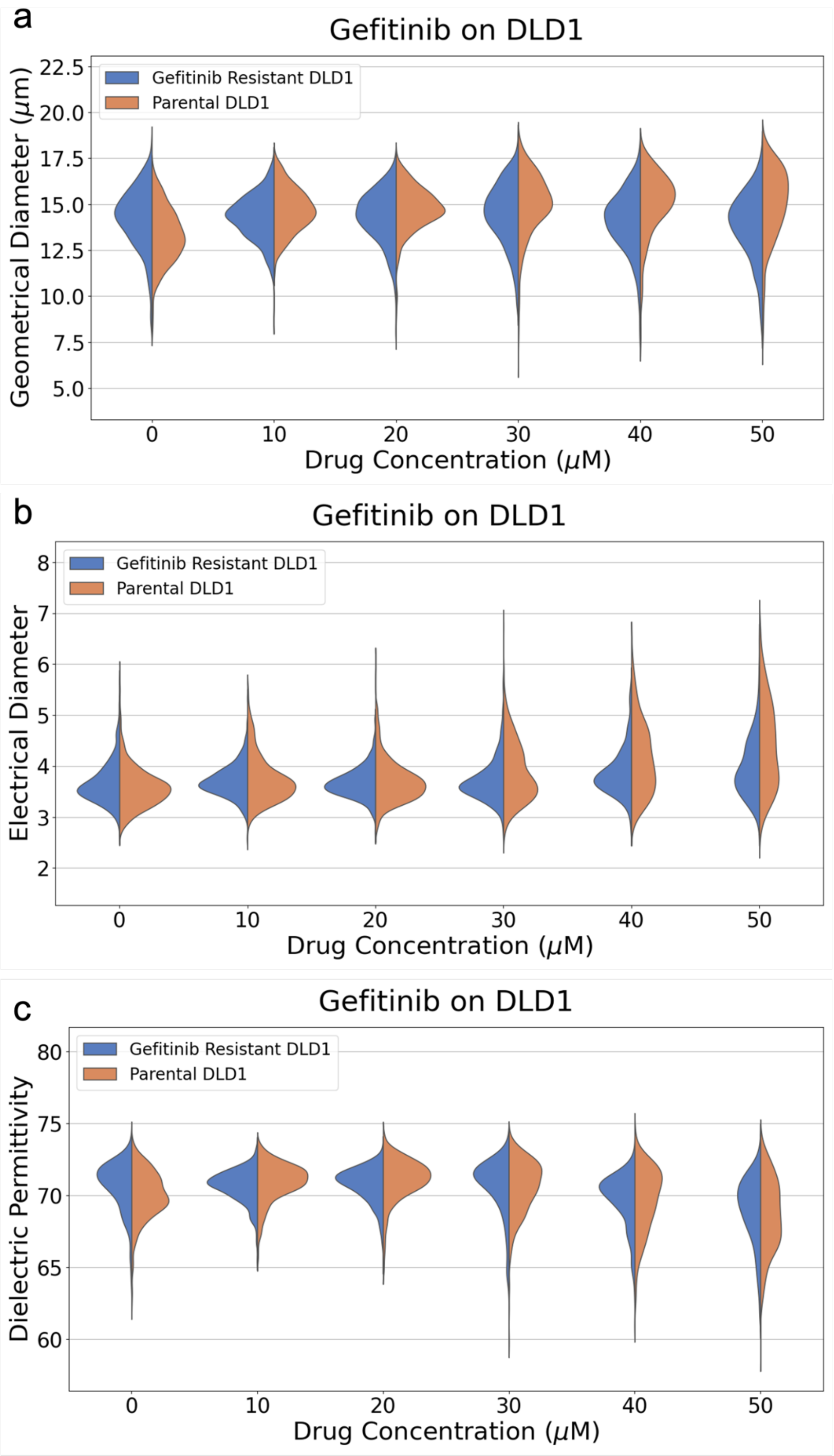

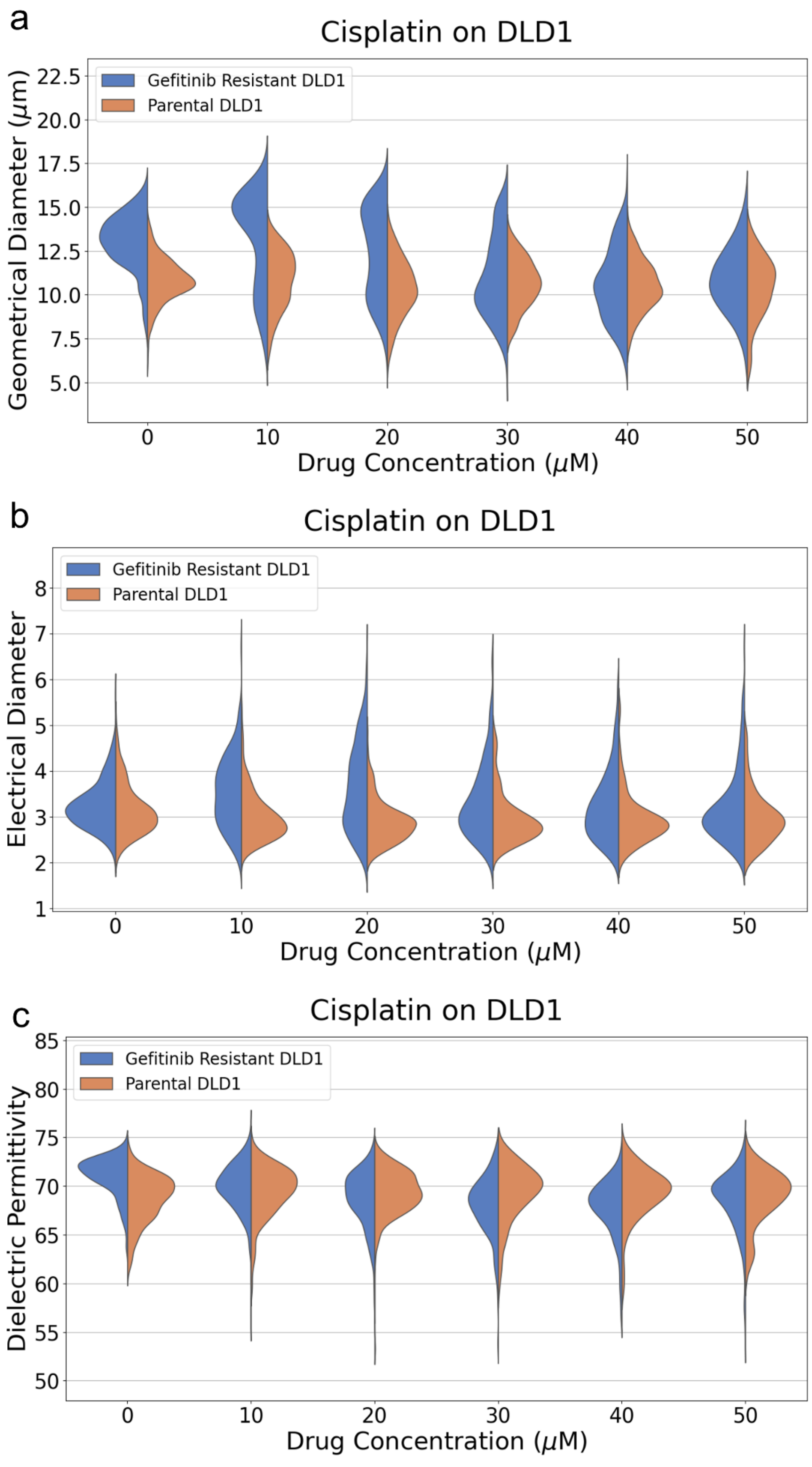
a
Cisplatin on DLD1
Gefitinib Resistant DLD1
Parental DLD1
Geometrical Diameter (μm)
22.5
20.0
17.5
15.0
12.5
10.0
7.5
5.0
0
10
20
30
40
50
Drug Concentration (μM)
b
Cisplatin on DLD1
Gefitinib Resistant DLD1
Parental DLD1
Electrical Diameter
8
7
6
5
4
3
2
1
0
10
20
30
40
50
Drug Concentration (μM)
c
Cisplatin on DLD1
Gefitinib Resistant DLD1
Parental DLD1
Dielectric Permittivity
85
80
75
70
65
60
55
50
0
10
20
30
40
50
Drug Concentration (μM)

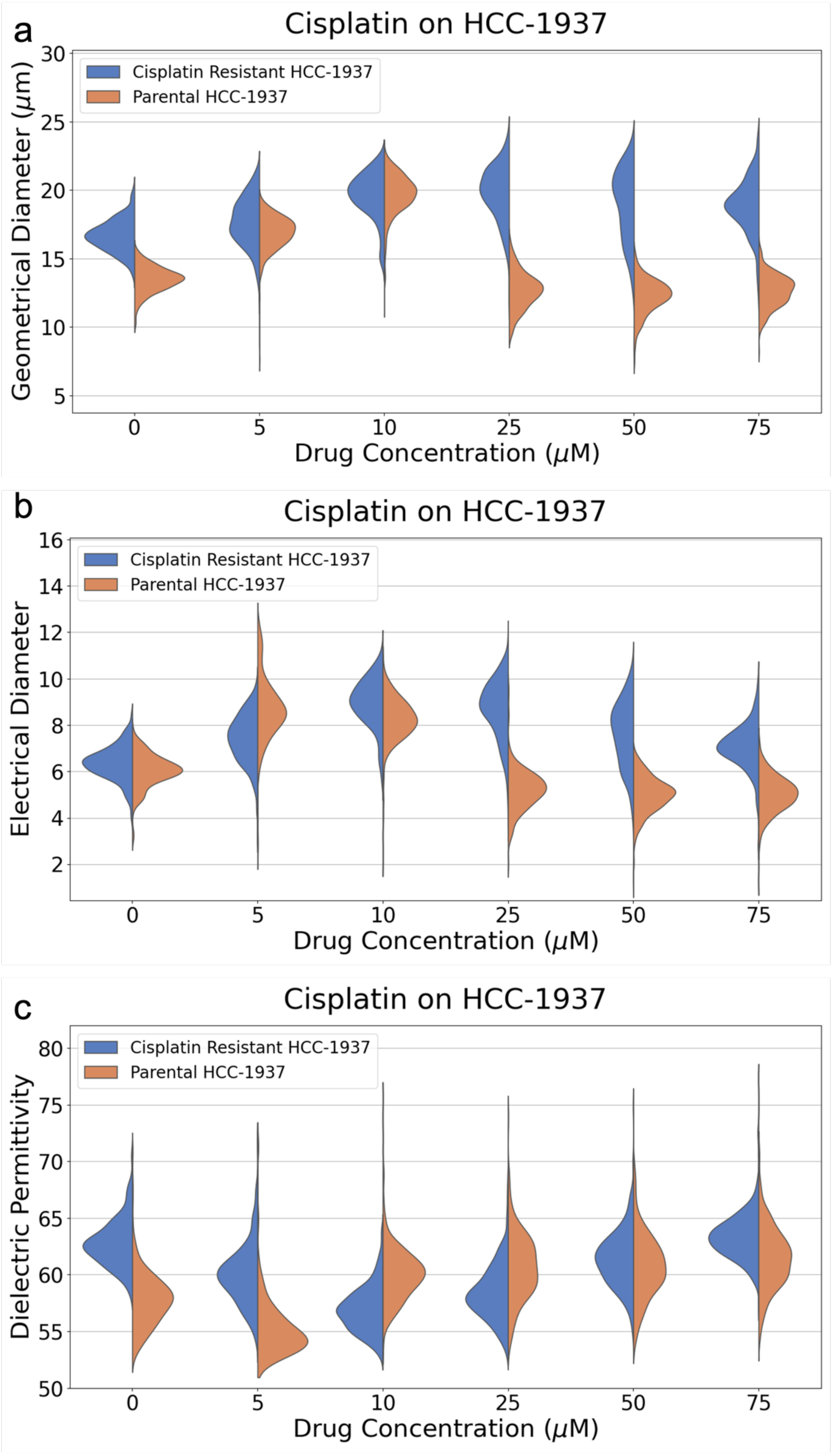
a
Cisplatin on HCC-1937
Cisplatin Resistant HCC-1937
Parental HCC-1937
Geometrical Diameter (μm)
30
25
20
15
10
5
0
5
10
25
50
75
Drug Concentration (μM)
b
Cisplatin on HCC-1937
Cisplatin Resistant HCC-1937
Parental HCC-1937
Electrical Diameter
16
14
12
10
8
6
4
2
0
5
10
25
50
75
Drug Concentration (μM)
c
Cisplatin on HCC-1937
Cisplatin Resistant HCC-1937
Parental HCC-1937
Dielectric Permittivity
80
75
70
65
60
55
50
0
5
10
25
50
75
Drug Concentration (μM)

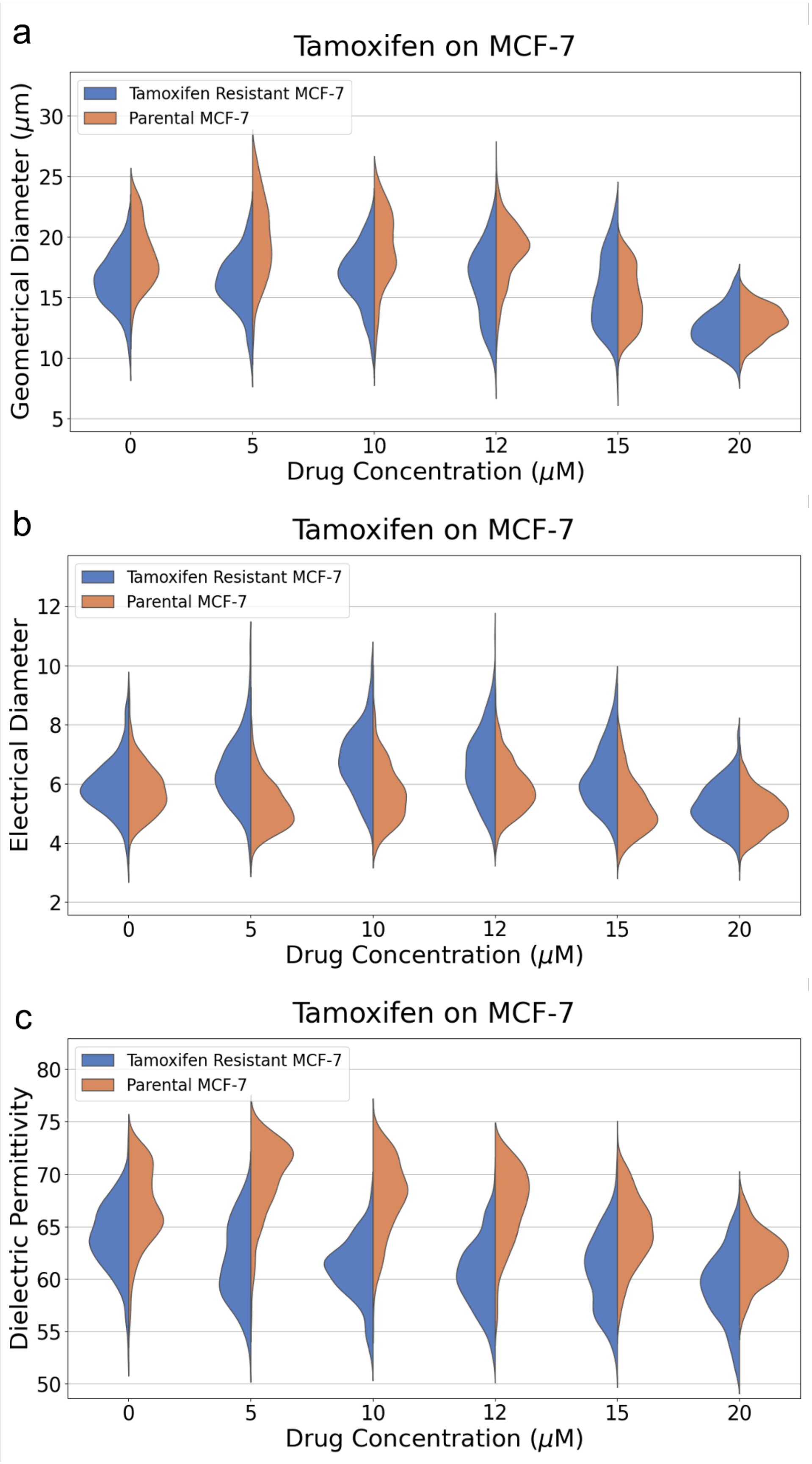
a
Tamoxifen on MCF-7
Tamoxifen Resistant MCF-7
Parental MCF-7
Geometrical Diameter (μm)
30
25
20
15
10
5
0
5
10
12
15
20
Drug Concentration (μM)
b
Tamoxifen on MCF-7
Tamoxifen Resistant MCF-7
Parental MCF-7
Electrical Diameter
12
10
8
6
4
2
0
5
10
12
15
20
Drug Concentration (μM)
c
Tamoxifen on MCF-7
Tamoxifen Resistant MCF-7
Parental MCF-7
Dielectric Permittivity
80
75
70
65
60
55
50
0
5
10
12
15
20
Drug Concentration (μM)

**Section 4. LC50 Graphs of the Various Cell Lines**

The following graphs shows the LC50 values indicating the development of drug resistance in the isogenic cell lines. The drug administrations were conducted in a 96-well plate. We note that the cell viability experiments reported in the main text were conducted in 6-well plates which might introduce variations in effective concentrations.

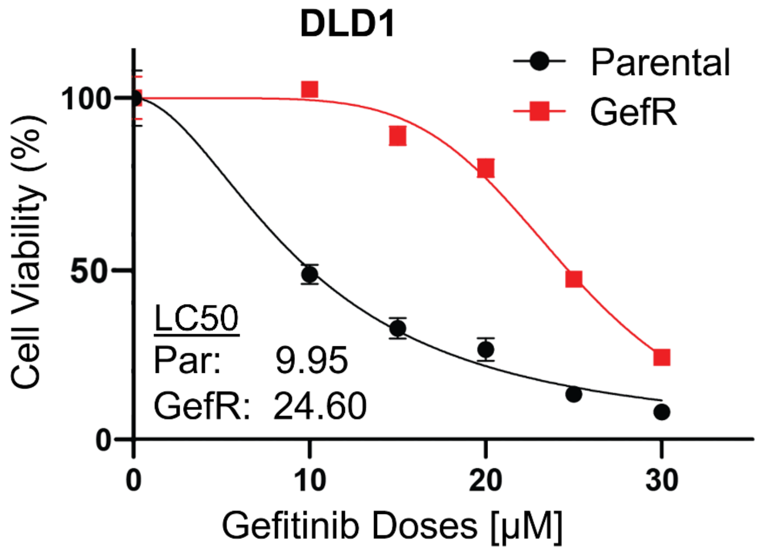


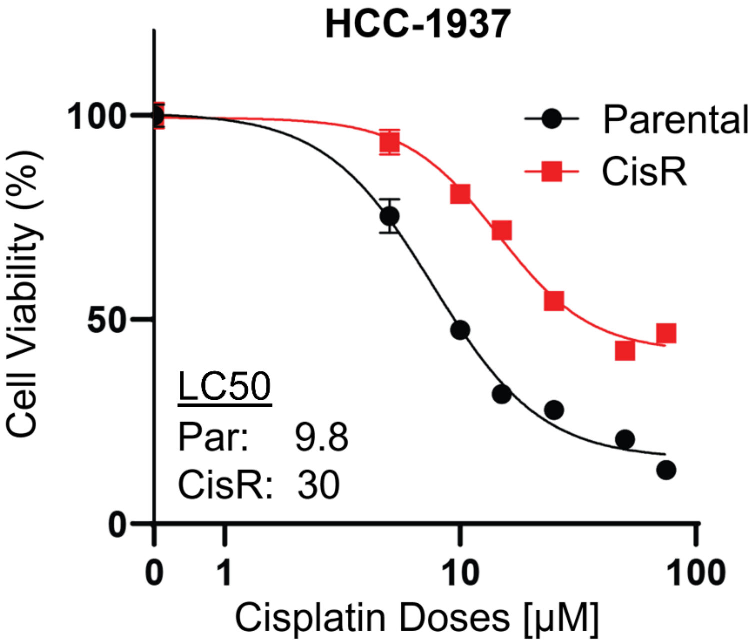

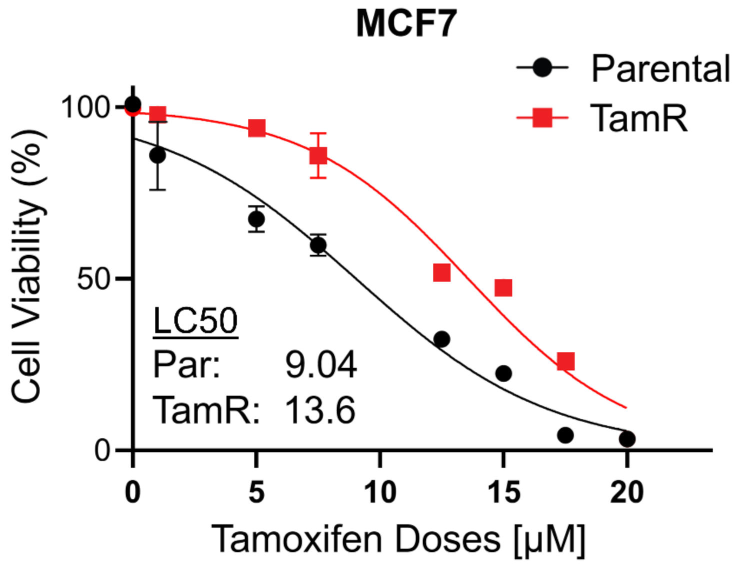


1 Coley, H. M., Shotton, C. F., Ajose-Adeogun, A., Modjtahedi, H. & Thomas, H. Receptor tyrosine kinase (RTK) inhibition is effective in chemosensitising EGFR-expressing drug resistant human ovarian cancer cell lines when used in combination with cytotoxic agents. *Biochemical pharmacology* **72**, 941-948 (2006).

2 La Monica, S. *et al.* Third generation EGFR inhibitor osimertinib combined with pemetrexed or cisplatin exerts long-lasting anti-tumor effect in EGFR-mutated pre-clinical models of NSCLC. *Journal of Experimental & Clinical Cancer Research* **38**, 1-12 (2019).

3 Ahsan, A. *et al.* Role of epidermal growth factor receptor degradation in cisplatin-induced cytotoxicity in head and neck cancer. *Cancer research* **70**, 2862-2869 (2010).
